# High-resolution crystal structure of a metabolic switch protein in a complex with monomeric c-di-GMP reveals a potential mechanism for c-di-GMP dimerization

**DOI:** 10.1101/2022.07.30.502141

**Authors:** Badri Nath Dubey, Viktoriya Shyp, Geoffrey Fucile, Urs Jenal, Tilman Schirmer

## Abstract

Bacterial second messengers c-di-GMP and (p)ppGpp have broad functional repertoires ranging from growth and cell cycle control to the regulation of biofilm formation and virulence. The recent identification of SmbA, an effector protein from *Caulobacter crescentus* that is jointly targeted by both signaling molecules, has opened up studies on how these global bacterial networks interact. C-di-GMP and (p)ppGpp compete for the same SmbA binding site, with a dimer of the former ligand inducing a conformational change of loop 7 leading to downstream signaling. Here, we report a crystal structure of a partial loop 7 deletion mutant, SmbA_Δloop_ in complex with c-di-GMP determined at 1.4 Å resolution. SmbA_Δloop_ binds monomeric c-di-GMP strengthening the view that loop 7 is required for c-di-GMP dimerization. In the crystal, SmbA_Δloop_ forms a 2-fold symmetric dimer via isologous interactions with the two symmetric halves of c-di-GMP. Structural comparisons of SmbA_Δloop_ with wild-type SmbA in complex with dimeric c-di-GMP or ppGpp support the idea that loop 7 is critical for SmbA function by interacting with downstream partners. These results underscore the flexibility of c-di-GMP in binding to the symmetric interface between protein subunits. It is envisaged that such isologous interactions of c-di-GMP will be observed in hitherto unrecognized targets.

## Introduction

In all domains of life, second messenger signaling is essential to modulate the intracellular response to external stimuli. In bacteria, purine nucleotide second messengers, such as guanosine tetra- and pentaphosphate, collectively referred to as (p)ppGpp, and bis-(3’-5’)-cyclic dimeric guanosine monophosphate, c-di-GMP, are involved in the global control of physiological responses to environmental change (Hauryliuk *et al*., 2015, Jenal *et al*., 2017;). (p)ppGpp is the primary regulator of bacterial growth and development in response to stress and nutrient limitation also known as the stringent response (Cashel & Gallant, 1969; Dalebroux & Swanson, 2012; Kalia *et al*., 2012). It modulates cellular reprogramming via multiple target proteins including RNA polymerase, translational GTPases, and metabolic enzymes, (Potrykus *et al*., 2011; Stott *et al*., 2015), thereby controlling bacterial transcription, translation (Wood *et al*., 2019), cell cycle progression (Gonzalez & Collier, 2014) (Lesley & Shapiro, 2008) stress resistance, and virulence (Martins *et al*., 2018; Nguyen *et al*., 2011). In most bacteria, c-di-GMP controls the transition between motile and sessile lifestyles. Low c-di-GMP levels are associated with motility, while its accumulation promotes adhesion and biofilm formation (Boehm *et al*., 2010; Baraquet & Harwood, 2013) (Fazli *et al*., 2014; Matsuyama *et al*., 2016). However, an increasing number of studies indicate that c-di-GMP has an impact on diverse aspects of bacterial physiology including cell cycle progression, metabolism, stress resistance, *etc*. (Morgan *et al*., 2014; Römling *et al*., 2005; Tschowri *et al*., 2014; Lori *et al*., 2015; Dubey *et al*., 2016, 2020)

The pleiotropic effects of (p)ppGpp and c-di-GMP are realized due to the diversity of their effectors, represented mainly by nucleotide-binding proteins and riboswitches (Li & He, 2012; Leduc & Roberts, 2009; Baraquet & Harwood, 2013). In particular, the structural diversity of the cyclic nucleotide, comprising various conformations from an extended monomeric form to a stacked dimer, explains the variety in c-di-GMP-binding motifs (Krasteva & Sondermann, 2017, Hengge, 2009, 2016; Chou & Galperin, 2016). The canonical c-di-GMP binding sites are represented by RxxxR and [DN]xSxxG motif in the PilZ domains, RxxD motif in degenerate GGDEF I site of DGCs and ExLxR in the EAL domains of PDEs. Moreover, several proteins with a non-canonical c-di-GMP binding motif have been recently characterized as high-affinity binding receptors, suggesting a widespread function of c-di-GMP in bacteria (Krasteva & Sondermann, 2017; Wang *et al*., 2016).

The development of biochemical methods to identify second messenger effectors greatly complemented our knowledge of novel c-di-GMP and/or (p)ppGpp binding proteins and their interaction networks (Nesper *et al*., 2012; Wang *et al*., 2019). Recently we have identified the first common target of c-di-GMP and ppGpp, SmbA protein from *C. crescentus* (Shyp *et al*., 2021). SmbA stimulates *Caulobacter* growth on glucose while preventing surface attachment in its active state repressed by binding of the c-di-GMP dimer (Fig 1). The two ligands inversely regulate protein activity presumably by affecting its conformation. The major conformational changes promoting SmbA functional switch affect the C-terminus helix 9 and the flexible loop 7 containing c-di-GMP subsite residues R211 and D214 from the RxxD motif (Fig 1). In the c-di-GMP-bound state, C-terminus helix 9 is stabilized by a salt bridge of D218 (from the loop7) and R289 (from helix 9), while in the ppGpp-bound state, loop 7 is disordered and helix 9 is in the open conformation (Fig 1). *In vivo* study revealed prolonged adaptation phases and reduced growth rates for the R211A mutant suggesting the involvement of loop 7 and potentially helix 9 in downstream signaling (Shyp *et al*., 2021).

**Figure 1.**
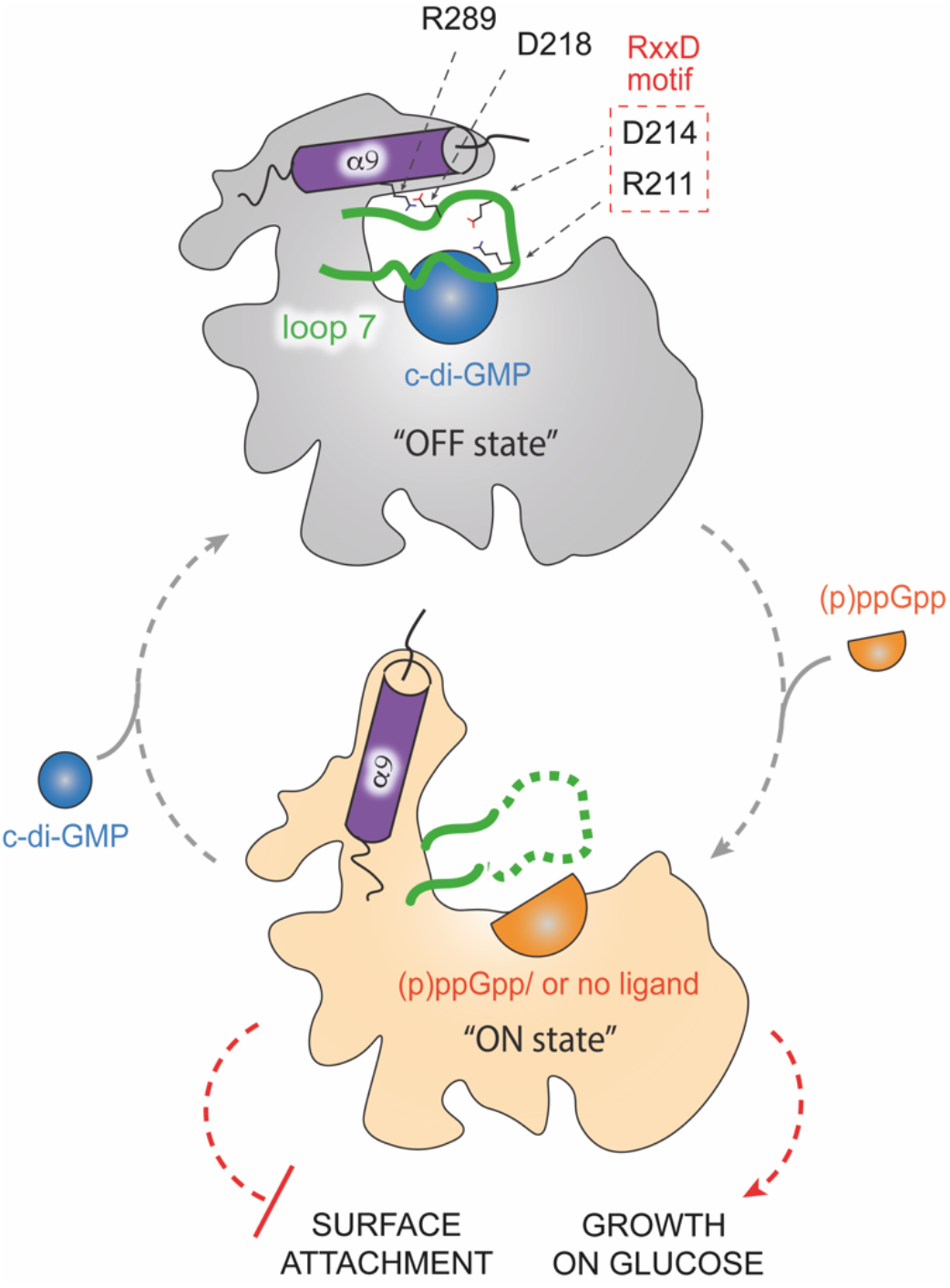
Second messenger mediated regulation of SmbA. Binding of a c-di-GMP dimer (blue sphere) inactivates SmbA (“OFF state”, grey), while its dissociation or displacement by a ppGpp monomer (an orange half-sphere) activates the protein (!ON state”, light orange). Loop 7 is shown in green, the C-terminal α9 helix is represented by a magenta cylinder. Amino acid residues essential for salt bridge formation between α9 helix and loop 7 are indicated. Key residues of the RxxD motif in loop 7 are shown in the red box. The physiological functions of activated SmbA are indicated with red dashed lines (Adopted from Shyp *et al*., 2021)).

To date, our structural knowledge about SmbA, however, is restricted to the wild-type protein in the presence of ligands. To understand how flexible loop7 influences the overall SmbA structure and its ligand binding, we present here the high-resolution structure of a loop 7 deletion mutant (fragment 198-215, hereafter SmbA_Δloop_). We observe that the mutant retains the TIM-barrel fold, however, accommodates only a monomer of c-di-GMP in a unique extended/open conformation. Importantly, in SmbA_Δloop_ mutant, C-terminal helix 9 adopts an outward orientation similar to that found in ppGpp-bound active state protein. Moreover, changes in c-di-GMP binding stoichiometry in SmbA_Δloop_ mutant, similar to loop 7 single mutant R211A, provide a potential mechanism and essential role of loop 7 in c-di-GMP dimerization and SmbA functional regulation.

## Results and discussion

### SmbA_Δloop_ forms a crystallographic dimer mediated by monomeric c-di-GMP

Ligand-induced conformational changes may be critical for SmbA physiological function, in particular for interaction with its yet-to-be-discovered downstream targets. Based on the fact that loop7 is disordered in the apo-state but becomes ordered upon binding of a c-di-GMP dimer, and that mutation of the interacting arginine residue 211 from this loop renders SmbA inactive in signaling (Shyp *et al*., 2021), we hypothesize that loop 7 is a central component of the physiological switch.

To explore the structural changes promoted by c-di-GMP via loop 7 we tried to crystallize the apo form of SmbA protein as well as SmbA_R211A_ and the SmbA_Δloop_ mutant with partial loop deletion (fragment 198-215) in complex with c-di-GMP. We only obtained suitable crystals for SmbA_Δloop_ (S1A Fig), which diffracted extremely well to 1.4 Å resolution and belong to space group P4_3_2_1_2 with one molecule in the asymmetric unit. The structure was determined by molecular replacement using the structure of wild-type SmbA (PDB: 6GS8) after removing c-di-GMP from the model as a template, followed by iterative refinement. The data collection and refinement statistics are summarized in Table 1.

**Table 1.** Crystallographic data collection and refinement statistics.

|  |  |
| --- | --- |
| Data collection |  |
| Synchrotron source | SLS, PXIII |
| Wavelength (Å) | 1.00004 |
| Space group | P 4<sub>3</sub> 2<sub>1</sub> 2 |
| a, b, c (Å) | 56.0, 56.0, 205.1 |
| α, β, γ (°) | 90, 90, 90 |
| Resolution (Å) | 21.5-1.4 (1.45-1.4) * |
| Unique reflections | 65577 (6381) |
| Completeness | 99.93 (99.87) |
| I/σ (I) | 20.6 (2.6) |
| Redundancy | 22.9 (22.9) |
| R<sub>merge</sub> (%) | 9.8 (164) |
| R<sub>pim</sub> (%) | 2.1 (352) |
| CC (1/2) % | 99.9 (86.4) |
| Refinement |  |
| R<sub>work</sub>/R<sub>free</sub> (%) | 14.8/17.7 |
| RMSD |  |
| Bond length (Å) | 0.006 |
| Bond angles (°) | 0.9 |
| Molecules/asymmetric unit | 1 |
| No. of atoms |  |
| Protein | 2081 |
| Ligand | 99 |
| Water | 299 |
| Average B-factor (Å<sup>2</sup>) | 20.4 |
| Protein | 18.5 |
| Ligand | 21.4 |
| Water | 33.3 |
| Ramachandran statistics (%) |  |
| Favored regions | 99.25 |
| Allowed regions | 0.75 |
| Disallowed regions | 0.0 |
| Deposition |  |
| PDB codes | 7B0E |

(\* = The values recorded in parentheses are those for the highest resolution shell)

The crystal structure shows that SmbA_Δloop_ forms a crystallographic dimer stabilized by a monomeric c-di-GMP molecule (Fig 2A). The ligand is found in a fully extended conformation and makes isologous interactions with the two protomers of the dimer (Fig 2B). The guanine bases of c-di-GMP interact extensively, via both polar and nonpolar contacts, with monomers A and B of the dimer. As in the wild-type complex, they form cation–π interactions with the guanidinium groups of R143 from both protomers (Fig 2B). Detailed interactions will be discussed in the next chapter. We have measured the c-di-GMP-to-protein stoichiometry using ITC. SmbA_Δloop_ yielded a K_D_ of 1.8 μM and a ligand-to-protein stoichiometry of 1:1 (S1B Fig), suggesting that c-di-GMP induced dimerization of SmbA_Δloop_ does not occur in solution and is due to the high protein concentration in the crystal.

**Figure 2.**
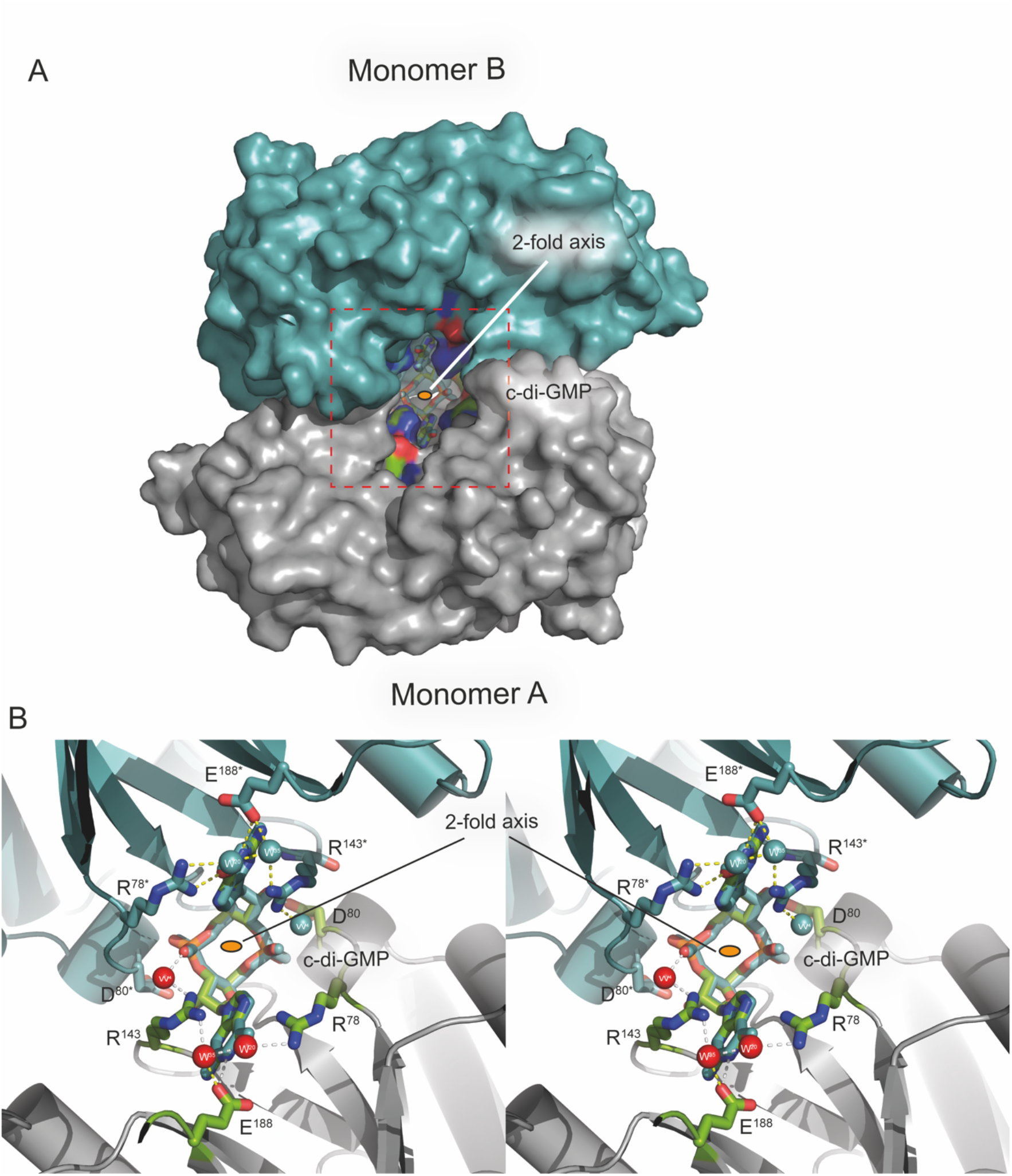
Crystal structure of SmbA_Δloop_ with c-di-GMP bound across crystallographic dyad. **(A)** The two monomers are depicted as surface (negatively charged atoms in red, positively charged atoms in blue and carbon atoms in green) with monomer A (gray) in standard orientation and monomer B (symmetry mate) in cyan. c-di-GMP (thick) in the dimer interface is shown as ball-and-stick model. **(B)** Stereoview down the twofold axis (indicated as a small orange ellipsoid), showing c-di-GMP forming isologous interactions with the two SmbA_Δloop_ protomers. Relevant residues are shown as color-coded sticks (oxygen, red; nitrogen, blue; carbon, green or cyan and waters as red and cyan spheres) and labeled. Residues and waters of the symmetry mate monomer are marked with an asterisk. Hydrogen bonds between subunits and c-di-GMP are indicated as yellow dotted lines.

C-di-GMP induced dimerization has been observed previously, involving monomeric, dimeric, and tetrameric c-di-GMP in the case of STING (Shang *et al*., 2012), VpsT (Krasteva *et al*., 2010), and BldD (Tschowri *et al*., 2014)), respectively (for a review see Chou & Galperin, 2016). Interestingly, structure comparison shows that the first two examples and SmbA involve symmetric stacking interactions, by R143 (SmbA), W131 (VpsT) and Tyr167 (STING) which cap two guanine bases of c-di-GMP from both sides at the dimer interface (S2 Fig). We anticipate that c-di-GMP mediated dimerization involving isologous interactions may be operational in hitherto unrecognized targets.

### Crystal structure of SmbA_Δloop_ with c-di-GMP and its comparison to the wild-type protein in complexes dimeric c-di-GMP and ppGpp

Overall, the SmbA_Δloop_ mutant retains the TIM-barrel fold with eight α-helices on the outside and eight parallel β-strands on the inside with an extra helix 9 (Fig 3A). The occupancy of the c-di-GMP ligand was set to 50% to account for its binding across the crystallographic dyad (half of the c-di-GMP molecule belongs to the symmetry mate). The ligand fit to the electron density very well after considerable conformational adjustment of both guanine bases (Fig 3B and S3 Fig). Thus, the mutant can accommodate only monomeric c-di-GMP, likely due to the absence of R211 and D214 of the RxxD motif of loop 7 essential for c-di-GMP dimer coordination (Fig. 3B).

**Figure 3.**
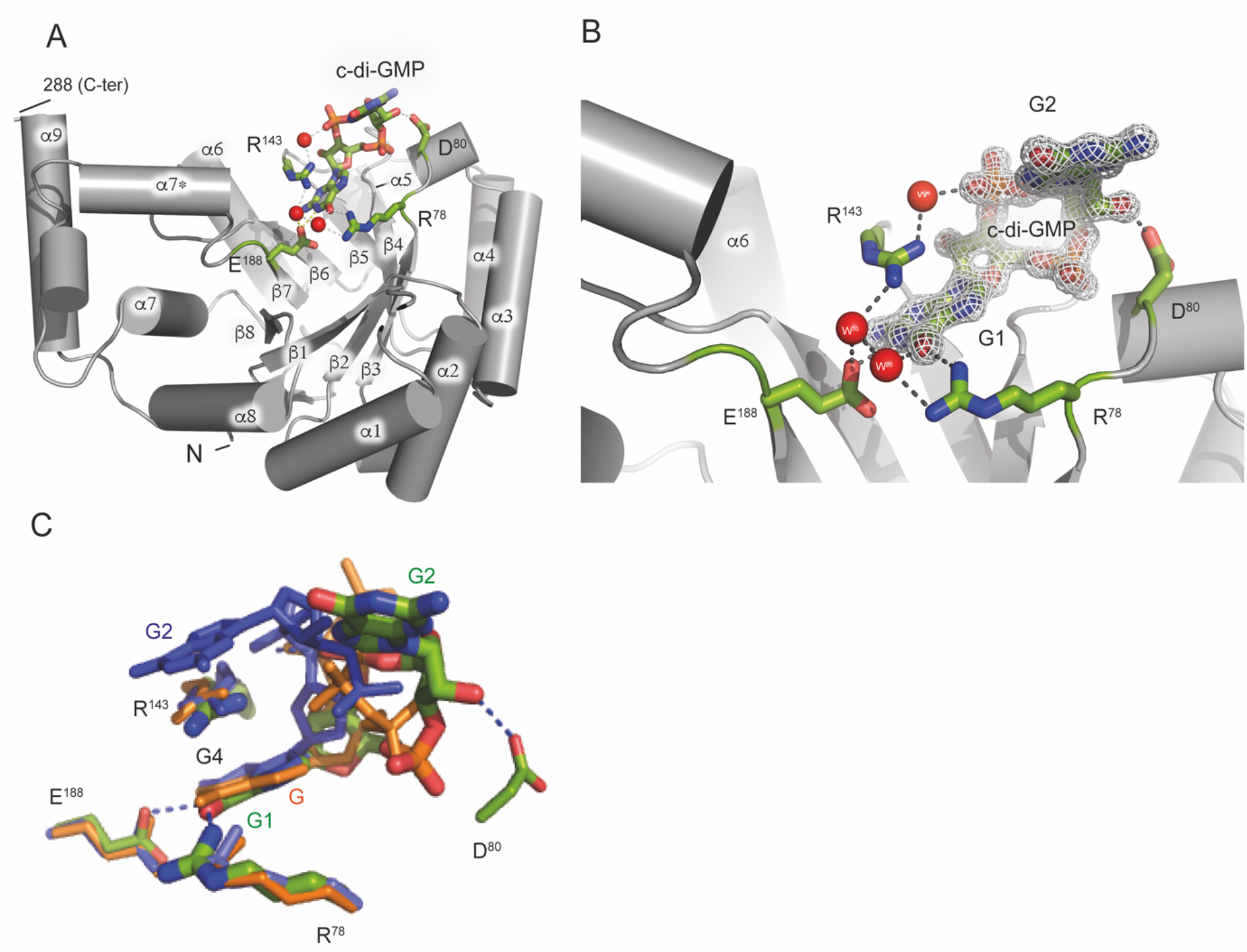
Electron density of c-di-GMP as bound to SmbA_Δloop_ and structural comparison with wild-type SmbA ligands. **(A)** Crystal structure of the SmbA_Δloop_ with the backbone drawn in grey cartoon and monomeric c-di-GMP shown in a stick. Residues in the SmbA_Δloop_ important in interaction with the c-di-GMP molecule are drawn in stick representation. Carbon atoms are shown in green, nitrogen in blue and oxygen in red. **(B)** 2Fo-Fc omit maps contoured at 1.2 σ of c-di-GMP and full structural details of the interacting residues. H-bonds (length < 3.5 Å) are indicated by gray lines and water molecules in red spheres. **(C)** View of c-di-GMP (green) as bound to SmbA_Δloop_, and the proximal c-di-GMP molecule (blue) of dimeric c-di-GMP and ppGpp (orange) as bound to wild-type SmbA_Δloop_. The proximal guanyl of monomeric c-di-GMP (G1), guanyl of ppGpp (G) and G4 of dimeric c-di-GMP overlap closely. While the other guanyl (G2) of the monomeric ligand has moved out considerably, to form isologous interactions with the second SmbA_Δloop_ molecule (not shown).

As shown above, the monomeric c-di-GMP ligand forms isologous interaction with the two protomers of the dimer (Fig. 2). The interactions of each guanyl with the protein are the same as observed for the proximal guanyl moiety (G4) of dimeric c-di-GMP and G of ppGpp interacting with wild-type SmbA (Shyp *et al*., 2021) (Fig. 3C). R143 is found stacked upon the guanyl to form a cation–π interaction, AR78 forms an H-bond with O6, and E188 forms on H-bond with N1 of the guanyl base (Fig. 3B, Fig S3). Compared to the wild-type complex the phosphate has moved towards the protein and forms an H-bond with main-chain amide 80 (Fig 3C). Three well-defined water molecules make hydrogen bonds with R78, E188, and R143 (Fig 3B).

Structural superimposition of SmbA_Δloop_/c-di-GMP with SmbA_wt_/(c-di-GMP)_2_ (6GS8) and SmbA_wt_/ppGpp (6GTM) shows RMS deviations of SmbA_Δloop_/c-di-GMP of 0.39 Å (for 214 Cα atoms) and 0.46 Å (for 230 Cα atoms) when compared to SmbA_wt_/c-di-GMP (Fig 4A) and SmbA/ppGpp, respectively (Fig 4B). These values show the high overall similarity of the structures accompanied by some notable local deviations. Particularly, in the SmbA_Δloop_/c-di-GMP complex, the C-terminal part of loop 7 forms a short helix α7* (Fig 3A). In addition, significant changes are observed in the C-terminal helix 9, which, in the wild-type protein, is stabilized by loop 7 being in turn immobilized by dimeric c-di-GMP. Thereby, the G1 and G2 guanyl bases interact with the RxxD motif of loop 7 (Shyp *et al*., 2021). In the SmbA_Δloop/_c-di-GMP complex, the monomeric ligand adopts an outward-open conformation similar to that found in SmbA_WT_/ppGpp complex (Fig 4B). However, its guanyl is in the same position as G of ppGpp and G4 of c-di-GMP all forming interactions with R78 and R143 (Fig 2C and Fig 4). At the same time, the phosphate moieties of monomeric c-di-GMP do not superimpose with those of bound dimeric c-di-GMP or ppGpp as bound to wild-type SmbA (Fig 2C). This observation further supports our model of the SmbA functional switch between on and off state upon ligand binding accompanied by structural changes in flexible loop 7 but not in the core structure.

**Figure 4.**
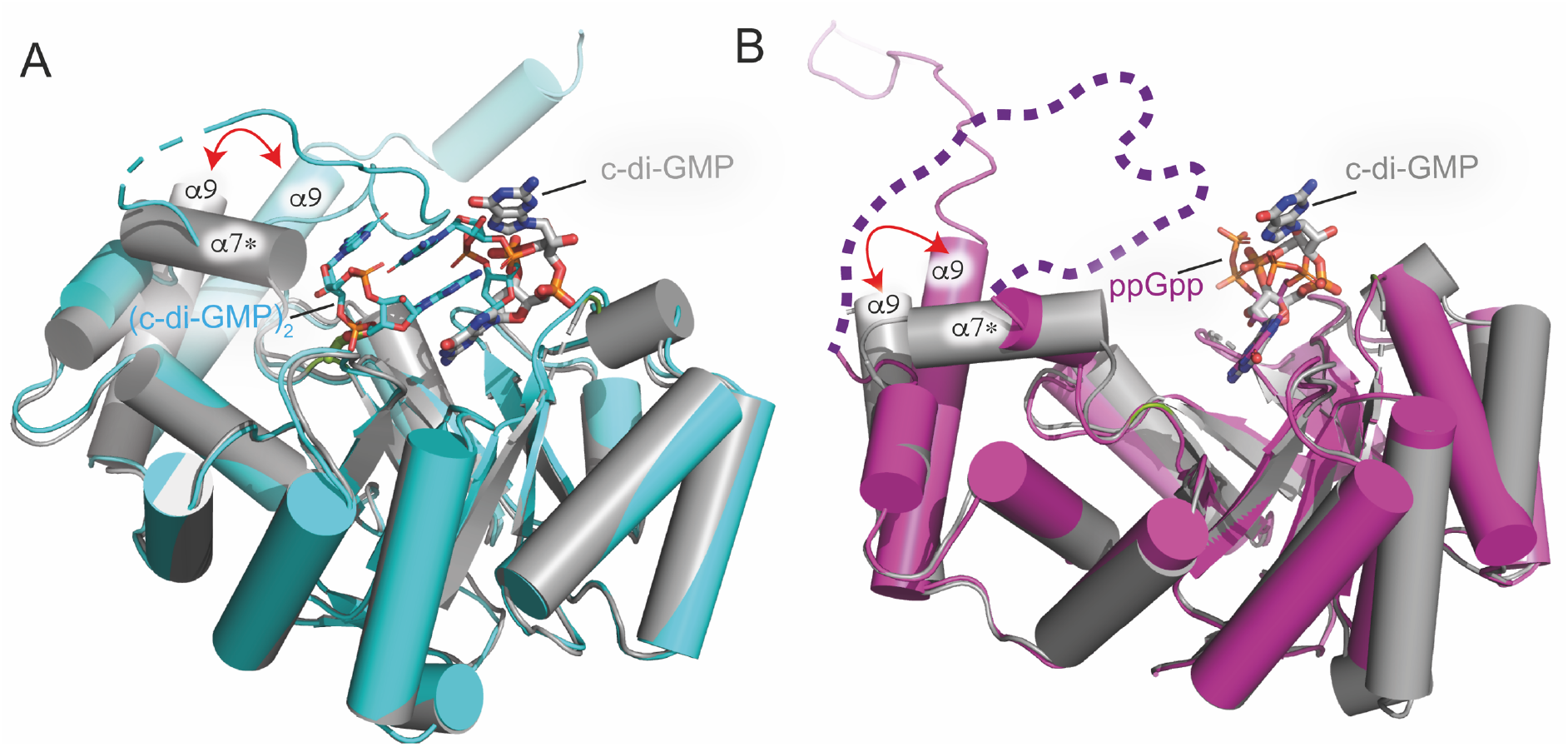
Structural comparison of SmbA_Δloop_/c-di-GMP with SmbA_wt_/c-di-GMP and SmbA_wt_/ppGpp. **(A)** Superposition of SmbA_Δloop_/c-di-GMP (gray) with SmbA_wt_/c-di-GMP (cyan) with RMSD of 0.4. Relevant secondary structure elements are labeled. Dimeric c-di-GMP (cyan) and monomeric (thick) are shown as ball-and-stick models. **(B)**Superposition of SmbA_Δloop_/c-di-GMP (gray) with SmbA/ppGpp (Magenta) with RMSD of 0.5. Relevant secondary-structure elements are labeled. ppGpp (magenta) and monomeric (thick in gray) are shown as ball-and-stick models. The disordered part of loop 7 is marked by broken lines.

### Conformation of the monomeric c-di-GMP bound to SmbA_Δ_loop

As discussed above, with the deletion of the loop containing the RxxD motif SmbA loses its ability to bind intercalated dimeric c-di-GMP molecule but still can hold one c-di-GMP. The monomeric ligand is two-fold symmetric, where the sugar pucker is C3′-endo, and both glycosidic torsion angles have a value of -126° (Figs 5A and 5B), which is significantly distinct to the *trans* conformation of G4 as part of dimeric c-di-GMP bound to wild-type SmbA. Superposition of c-di-GMP from the SmbA_Δloop_ and SmbA_wt_ complex structures shows that this difference is the reason for the elongated shape of monomeric c-di-GMP, while macrocycle including the sugar superimposes closely (Fig 5C).

**Figure 5.**
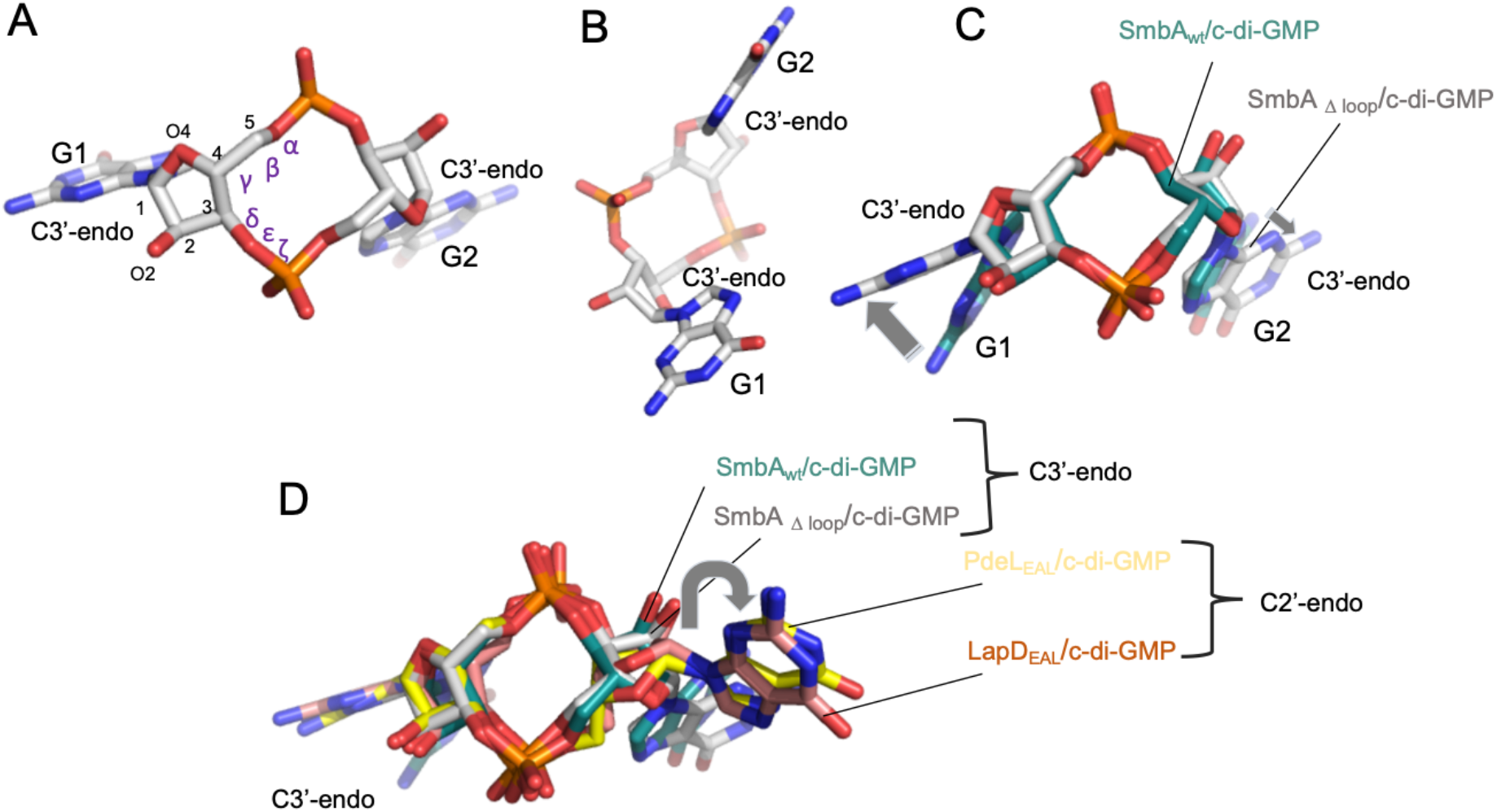
Observed c-di-GMP conformations in SmbA_loop_ and its comparison with SmbA_wt_, PdeL and LapD. **A)** and **(B)** shows the partial open-twisted form of monomeric c-di-GMP in C3’-endo sugar pucker conformation observed in SmbA mutant. **(C)** Superimposition of c-di-GMP from SmbA_Δloop_. GMP moiety from both structures shows the same conformation, the C3′-endo sugar pucker; however, there are considerable differences in the G1 and G2 base orientation (indicated by the gray arrow). **(D)**Superimposition of c-di-GMP from SmbA_Δloop_ with monomeric c-di-GMP as observed when bound to a phosphodiesterase PdeL and degenerated-phosphodiesterase LapD. Distinct sugar pucker of the base at the right (G2) appears responsible for the fully elongated form of c-di-GMP when bound to PdeL or LapD. In contrast, all bases at the left (G1) show the same supar pucker, i.e. C3′-endo as also observed for SmbA_Δloop_ in this study.

Next, we were interested in comparing the conformation of monomeric c-di-GMP as bound to SmbA_**Δloop**_ to other effectors that bind the ligand in the monomeric form such as the phosphodiesterase domain PdeL_EAL_ (Sundriyal *et al*., 2014) and the degenerate LapD_EAL_ domain (Navarro *et al*., 2011). A superposition of the three complexes is shown in Fig 5D. While the macrocycles retain a similar, but not identical, conformation, as seen in the SmbA_Δloop,_ the ligands bound to PdeL_EAL_ and LapD_EAL,_ are in a more open conformation, apparently due to the C2′-endo puckering of one of the guanines (at the right side in Fig 5D). These results show that c-di-GMP can adopt yet another unique conformation different from the stacked dimeric conformation in complex with SmbA_wt,_ or the extended form in the PdeL_EAL_ (PDB code-4LJ3) or degenerate LapD_EAL_ domain (PDB code-3PJT).

The monomeric c-di-GMP conformation observed in the SmbA_Δloop_ complex structure is different from that of dimeric c-di-GMP. From this comparison, one can see that one G1 is bound always the same way in the three complexes (Fig 3C). Due to the conformational changes, the other GMP has moved out considerably, to form an isologous interaction with the second SmbA_Δloop_ molecule (not shown) (Fig 3C). This indicates that, depending on its binding partner, c-di-GMP is flexible enough to adopt various conformations via only minor changes in torsion-angle.

### Phylogenetic analysis and exploring SmbA homologs

To understand the evolutionary significance of the flexible loop of SmbA switch protein, here we have further extended our primary sequence analysis of SmbA and its homologs described briefly in Shyp et al. 2021. We identified SmbA orthologs based on reciprocal best BLAST hits across species, concordance of the protein sequence distance tree with a species phylogeny based on 16S rRNA markers [Abraham et al., 2008] and syntenic conservation [Altenhoff et al., 2015] (Figs 6A and 6B). Interestingly, the c-di-GMP-binding RxxD motif is only strictly conserved within the *Caulobacter* genus, with either Asn or Glu substitution among the *Caulobacterales* (Fig 6C). There is considerable variability around this loop region, including several insertions and deletion events. This may suggest alternative binding modes and/or substrates within the *Caulobacterales* order. Similarly, the sites interacting with ppGpp (R78, N111, Q114, R143, E188) are not strictly conserved within the *Caulobacterales* order. The C-terminal helix 9 is highly conserved among SmbA orthologs (Fig 6C). This is consistent with the proposal that it adopts a different conformation in the c-di-GMP-bound state than apo and ppGpp, thus necessary for the ligand-mediated SmbA switch. The strictly conserved N-terminal motif (MRYRP[FL]G) is also found in otherwise unrelated proteins from the *Acetomycetalesorder* (Frankia, Streptomyces).

**Figure 6.**
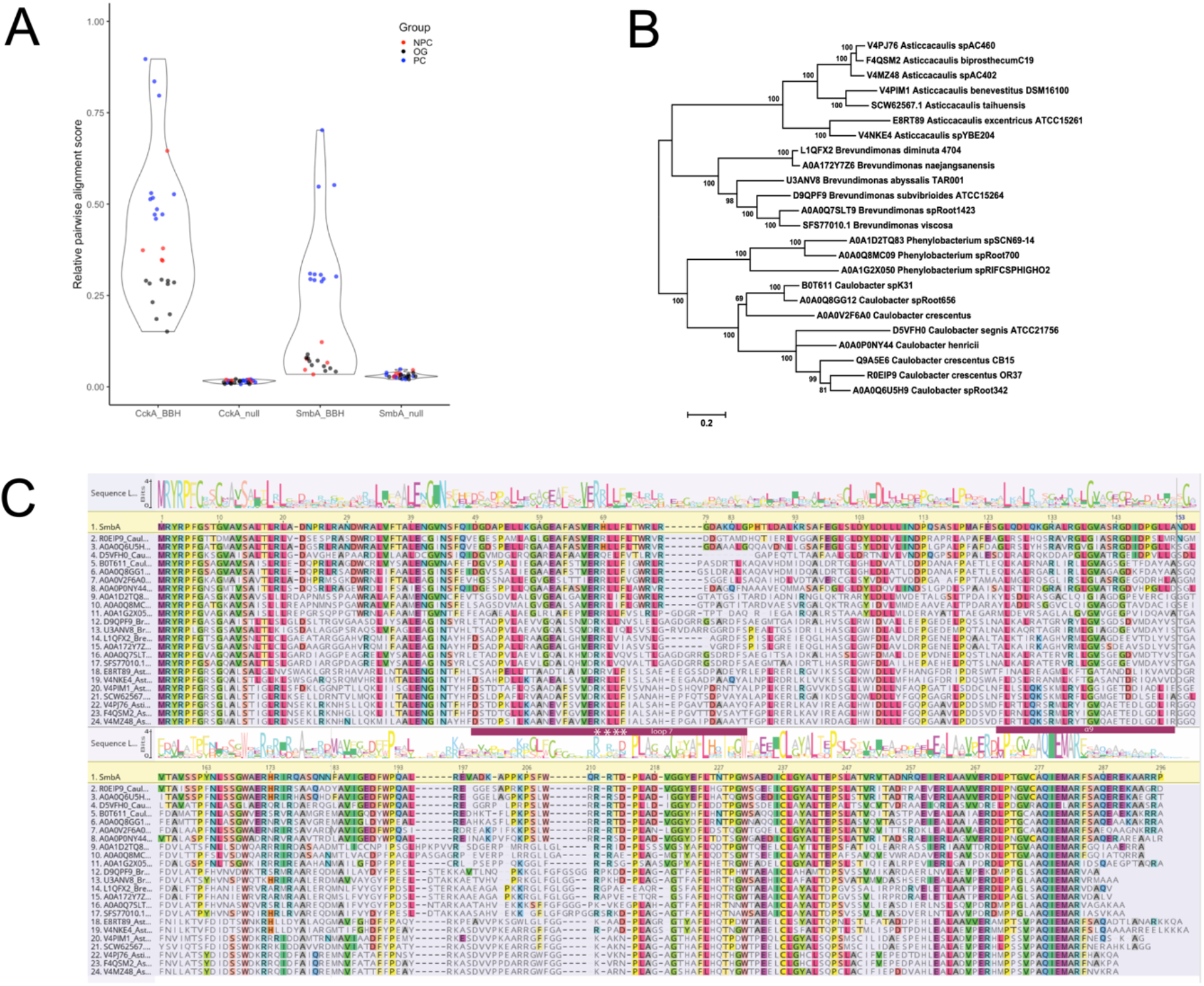
Sequence alignment and distance of SmbA homologs. **(A)** Pairwise Needleman-Wunsch global alignment scores of SmbA and CckA reciprocal best BLAST hits (BBH) for species sampled from prosthecate Caulobacterales (PC), non-prosthecate Caulobacterales (NPC), and other bacterial groups (OG). Alignment scores are reported relative to self-alignment of SmbA (Q9A5E6) and CckA (H7C7G9) from Caulobacter crescentus. For the null models, CckA BBH was scored against SmbA and vice versa.. The latter BBH was identified using BLASTp against the NCBI-NR database using the BLOSUM45 scoring matrix. **(B)** A phylogenetic tree of 24 SmbA orthologs inferred using the Maximum Likelihood method based on the JTT model as implemented in MEGA7. Branch lengths indicate the number of substitutions per site. The tree with the highest log likelihood (-9134.38) is shown, with bootstrap support from 100 replicates indicated at branches. **(C)** Sequence alignment and logo of SmbA orthologs. The sequence logo was generated using the WebLogo server from the global alignment of SmbA orthologs used to build the distance tree.

The *Caulobacterales* order contains prosthecate and non-prosthecate species (Abraham *et al*., 2008, 2014). SmbA (Q9A5E6) appears to be unique to the prosthecate *Caulobacterales*. Reciprocal best BLAST hits for SmbA from non-prosthecate *Caulobacterales* and other bacterial species are very distantly related Aldo-keto reductases which cannot be meaningfully aligned with SmbA. The central function of SmbA is a simple molecular switch that responds to the cellular concentrations of ppGpp and c-di-GMP to regulate *Caulobacter* growth (Shyp *et al*., 2021). We surmise that the presence of a SmbA ortholog is a marker for prosthecate-type *Caulobacterales* species which have not been morphologically characterized. This is further supported by the genes flanking SmbA, including a putative iron-sulfur glutaredoxin (Q9A5E5) and a BolA/YrbA family transcription factor (Q9A5E7) which in *E. coli* positively regulates the transition from the planktonic to attachment stage of biofilm formation (Dressaire *et al*., 2015)

## Material and methods

### Plasmid construction and purification of the recombinant protein

To construct pET21b-smbA_Δloop_-His6 (deletion of fragment 198-215), the pET21b-smbA-His6 plasmid was amplified with the following primers: 6265_D Loop7_forward CCCCAGGCCCTGCGAGAACTGGCCGATGTGGGCGGCTA and 6266_DLoop7_reverse TAGCCGCCCACATCGGCCAGTTCTCGCAGGGCCTGGGG. The template was digested with DpnI and mutant DNA was transformed into competent cells for nick repair. The final construct has been sequenced to confirm the fragment deletion. Protein was overproduced and purified as described previously (Shyp *et al*., 2021). Briefly, *E. coli* Rosetta 2(DE3) cells were used to overproduce recombinant protein from the pET21b expression plasmid. Cells were grown in LB-Miller supplemented with 100 μg/ml ampicillin to an OD_600_ of 0.4 to 0.6, expression was induced with 1 mM IPTG overnight at 22°C. Cells were harvested by centrifugation (5000 g, 20 min, 4°C), washed with PBS and flash-frozen in liquid N_2_, and stored at -80°C until purification.

For purification, cells were resuspended in *lysis buffer* (30 mM Tris/HCl pH 7.5, 5 mM MgCl_2_, 100 mM NaCl, 1mM DTT and 10 mM imidazole containing 0.2 mg/ml lysozyme, DNaseI (AppliChem) and Complete Protease inhibitor (Roche) and disrupted using a French press. The suspension was clarified by centrifugation at 30,000 x g (Sorval SLA 1500) at 4 °C for 30 min and loaded onto a 1 ml HisTrap HP column (GE Healthcare) on an ÄKTA purifier 10 system (GE Healthcare). Column was washed with 5 column volumes with *wash buffer* (30 mM Tris/HCl pH 7.5, 5 mM MgCl_2_, 100 mM NaCl, 1mM DTT and 10 mM imidazole), and the bound protein was eluted with linear gradient of *elution buffer* (30 mM Tris/HCl pH 7.5, 3mM MgCl_2_, 100 mM NaCl, 1mM DTT and 300 mM imidazole). Elution fractions enriched in SmbA (as judged by SDS-PAGE) were pooled and concentrated to around 10 mg/ml using Amicon Ultra centrifugal concentrator with a nominal molecular weight cut-off of 30 kDa (Millipore AG). The concentrated protein was centrifuged at 16,000 x g at 4°C for 15 min and loaded onto a Superdex 75 gel filtration column (Amersham Biosciences) equilibrated with 30 mM Tris/HCl pH 7.5, 5 mM MgCl_2_, 100 mM NaCl, 1mM DTT. Fractions containing essentially pure SmbA (as judged by SDS-PAGE) were pooled and concentrated to a desired concertation for further experiments.

### Crystallization

A Phoenix robot (Art Robbins Instruments) was used for a wide range of crystallization screening. Crystallization was carried out using the sitting drop vapour diffusion method at 20 °C by mixing the protein with the reservoir solution in a 1:1 ratio. The protein concentration was 5.0, 2.25 and 1.75 mg/ml upon adding c-di-GMP in 3.0 fold molar excess. Triangle diamond-shaped 3D crystals appeared in Pact premier D11 (Molecular dimension) after one week in 0.2 M Calcium chloride dihydrate 0.1 M Tris pH 8.0 and 20 % w/v PEG 6000. Crystals were flash-frozen into two different cryoprotectants. The best diffraction was obtained from crystals cryo-protected with 25% ethylene glycol.

### X-Ray diffraction data collection, phasing, and refinement

All single-crystal X-ray diffraction data sets were collected at PXIII beamline of Swiss Light source, Villigen, Switzerland.) Dataset was collected for the crystal of the SmbAΔloop in presence of c-di-GMP. Diffraction data sets were processed either with MOSFLM (Leslie, 2006) or XDS (Kabsch, 2010), and the resulting intensities were scaled using SCALA from CCP4/CCP4i2 suite(Potterton *et al*., 2018) For solving the SmbAΔloop structure, the refined SmbAwt (PDB code, 6GS8) structure was used as search model without c-di-GMP. The structure was solved by molecular replacement using PHENIX PHASER (McCoy *et al*., 2007) Further refinement of structures was carried out using REFMAC5 and Phenix refinement (Terwilliger *et al*., 2008). Model building was performed using COOT (Emsley & Cowtan, 2004) and model validation was carried out with molprobity (Williams *et al*., 2018). Crystallographic data processing and refinement statistics are provided in Table 1.

### Isothermal titration calorimetry (ITC)

Experiments were carried out at 25°C or 10°C, a syringe stirring speed of 300 rpm, a pre-injection delay of 200 secs, and a recording interval of 250 secs in a Microcal VP-ITC in ITC buffer (30 mM Tris-HCl pH 7.5, 150 mM NaCl, 5 mM MgCl_2_). All solutions were degassed below the temperature used in the experiments before loading into the calorimeter cell. Baseline correction and integration of the raw differential power data, and fitting of the resulting binding isotherms to obtain dissociation constants were performed using the Microcal ORIGIN software.

### Bioinformatics

BLAST analyses were conducted using the NCBI-NR dataset. Multiple sequence alignments were generated using MAFFT in G-INS-i mode (Katoh *et al*., 2005) followed by manual refinement. The phylogenetic tree of 24 SmbA orthologs was inferred using the Maximum Likelihood method based on the JTT model(Jones *et al*., 1992) as implemented in MEGA7 (Kumar *et al*., 2016). Branch lengths indicate the number of substitutions per site. The tree with the highest log likelihood (-9134.38) is shown, with bootstrap support from 100 replicates indicated at branches. Initial tree(s) for the heuristic search were obtained automatically by applying Neighbor-Join and BioNJ algorithms to a matrix of pairwise distances estimated using a JTT model, and then selecting the topology with a superior log-likelihood value. A discrete Gamma distribution was used to model evolutionary rate differences among sites (5 categories, +G = 2.2328)). The rate variation model allowed for some sites to be evolutionarily invariable ([+I], 7.16% sites). The tree is drawn to scale, with branch lengths measured in the number of substitutions per site. There were a total of 323 positions in the final dataset.

### Protein Data Bank deposition

The final SmbA_Δloop_ coordinates and structure factor amplitudes have been deposited in the Protein Data Bank and are available under accession number 7B0E.

## Author contributions

B.N.D. performed purifications, crystallization, and biophysical experiments. B.N.D. processed X-ray data, determined structure, built and validated the model. G.F. performed bioinformatics analysis. V.S. performed cloning. B.N.D and V.S. wrote the manuscript with input from all authors. B.N.D. V.S. U.J. and T.S. conceived and directed the project. This work was supported by the European Research Council (ERC) Advanced Research Grant (3222809) and the Swiss National Science Foundation (310030B_147090) to U.J and the Swiss National Science Foundation (grant no.31003A_166652) to T.S.

## Acknowledgments

We thank the beamline staff at the Swiss Light Source in Villigen for expert help in data acquisition and T. Sharpe (Biophysics Facility, Biozentrum, University of Basel) for excellent support. Bioinformatics analyses were performed at sciCORE (http://scicore.unibas.ch/) scientific computing center at the University of Basel, with support from the SIB - Swiss Institute of Bioinformatics.

## Supplementary figures

**S1 Fig.**
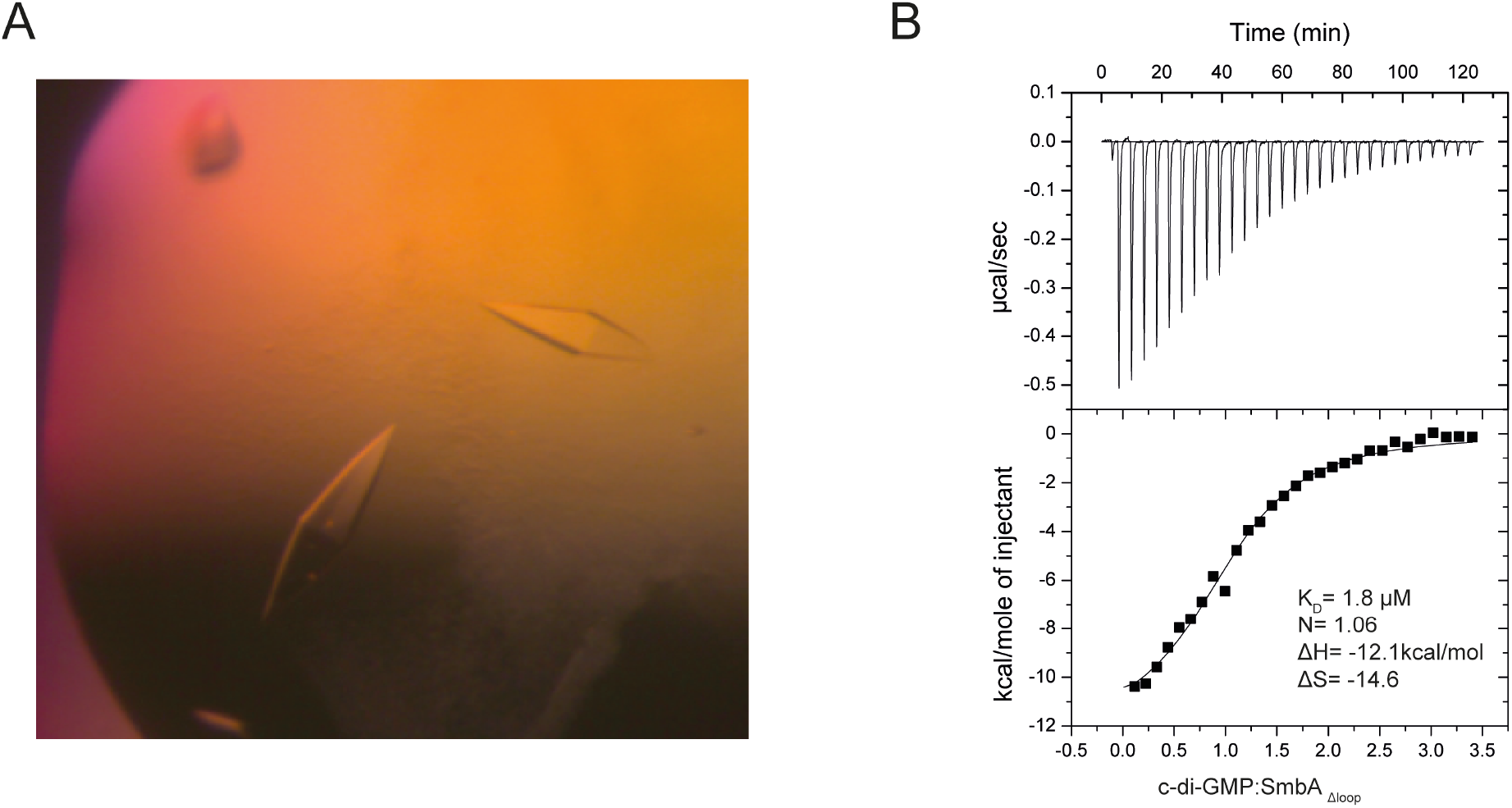
SmbA_Δloop_ crystals and binding parameters of SmbA_Δloop_ with c-di-GMP. **(A)** SmbA_Δloop_ crystals in complex with c-di-GMP. **(B)** ITC of SmbA_Δloop_ (10 μM) binding to the c-di-GMP molecule (150 μM). The binding stoichiometry, ΔH, ΔS, and K_D_ are marked.

**S2 Fig.**
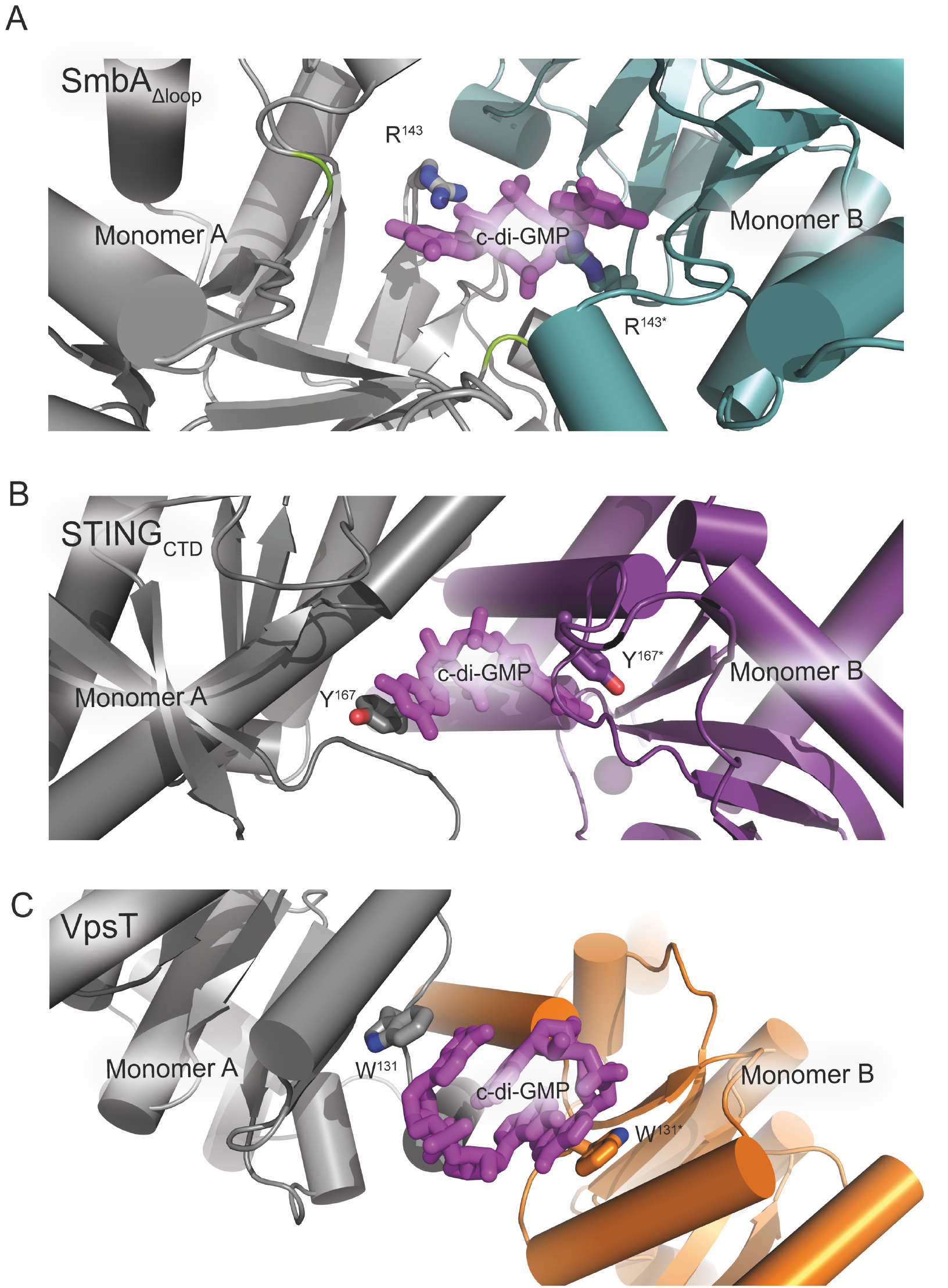
c-di-GMP bound at dimer interface found in STING and VpsT. c-di-GMP bind at the crystallographic dimer interface shown in the stick model (magenta). Residues involved in base stacking are shown in a stick. Monomers are colored differently. (A) SmbA_Δloop_ c-di-GMP contact at the 2-fold crystallographic dimer interface, (B) STING dimerization interface (PDB code-4F5Y). (C) VpsT dimerization interface (PDB code-3KLO).

**S3 Fig.**
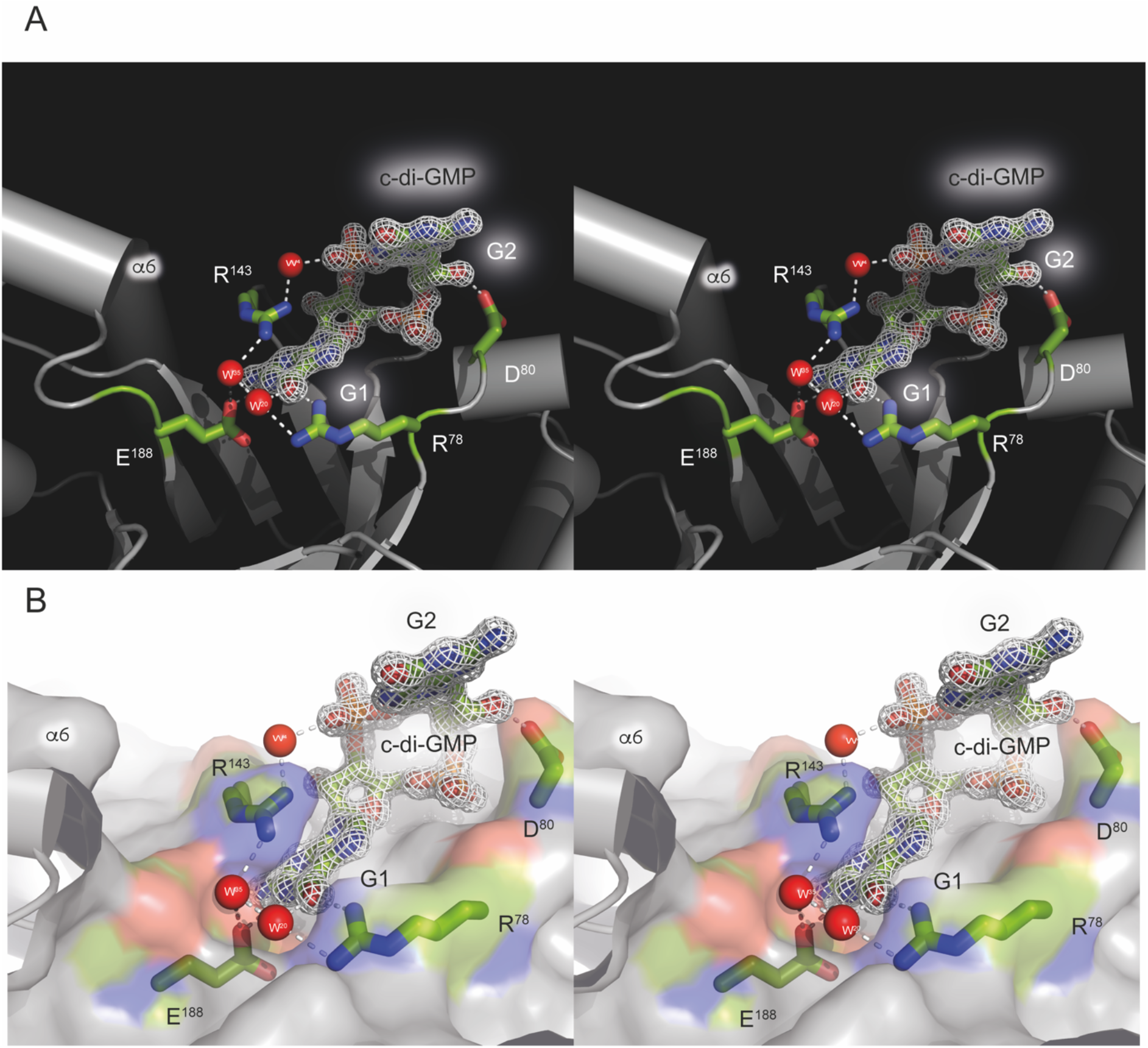
An expanded stereo view of the SmbA_Δloop_ residues interacting with c-di-GMP. **(A)** Stereo view of the 2Fo-Fc omit maps contoured at 1.2 σ. The molecular structure of c-di-GMP is embedded in the map. The colour code is similar to that in Fig. 3A. The R143 guanidinium group stacks very well with the guanyl base of c-di-GMP., while R78 is engaged in lateral H-bonding. **(B)** Stereo view of the c-di-GMP molecule drawn as surface representation (negatively charged atoms in red, positively charged atoms in blue and carbon atoms in green). Hydrogen bonds are marked by dotted lines in white.

