## Supplementary Information for "High-resolution crystal structure of a metabolic switch protein in a complex with monomeric c-di-GMP reveals a potential mechanism for c-di-GMP dimerization"

### Supplementary figures

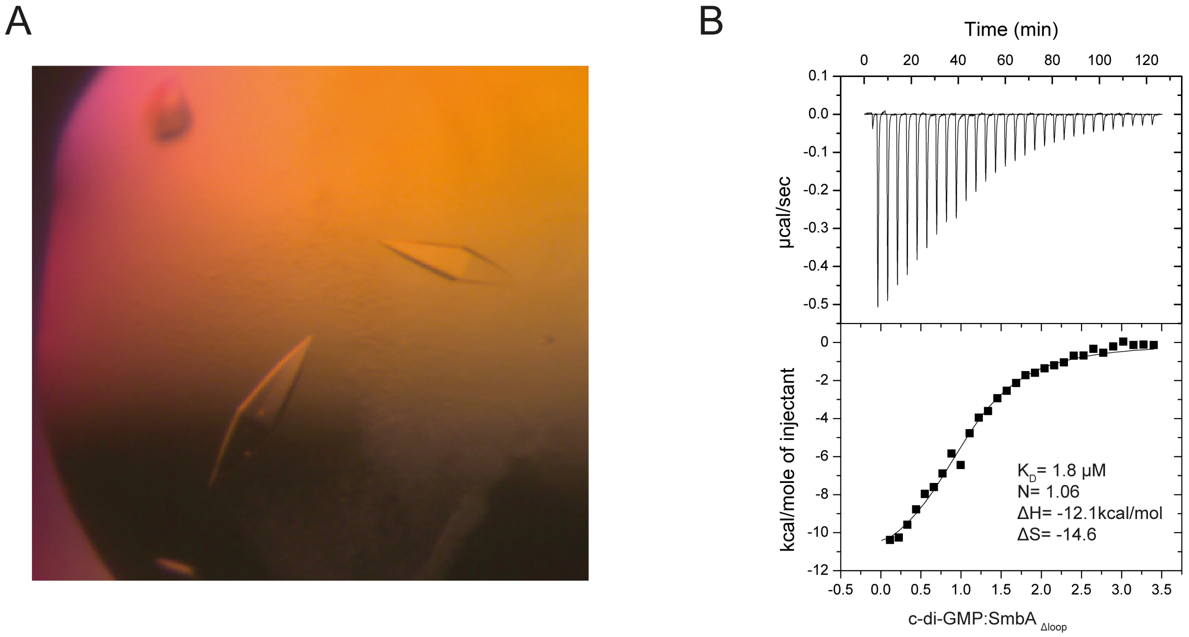


**S1 Fig. SmbA$_{\Delta loop}$ crystals and binding parameters of SmbA$_{\Delta loop}$ with c-di-GMP.**

**(A)** SmbA$_{\Delta loop}$ crystals in complex with c-di-GMP.

**(B)** ITC of SmbA$_{\Delta loop}$ (10 µM) binding to the c-di-GMP molecule (150 µM). The binding stoichiometry, ΔH, ΔS, and $K_D$ are marked.

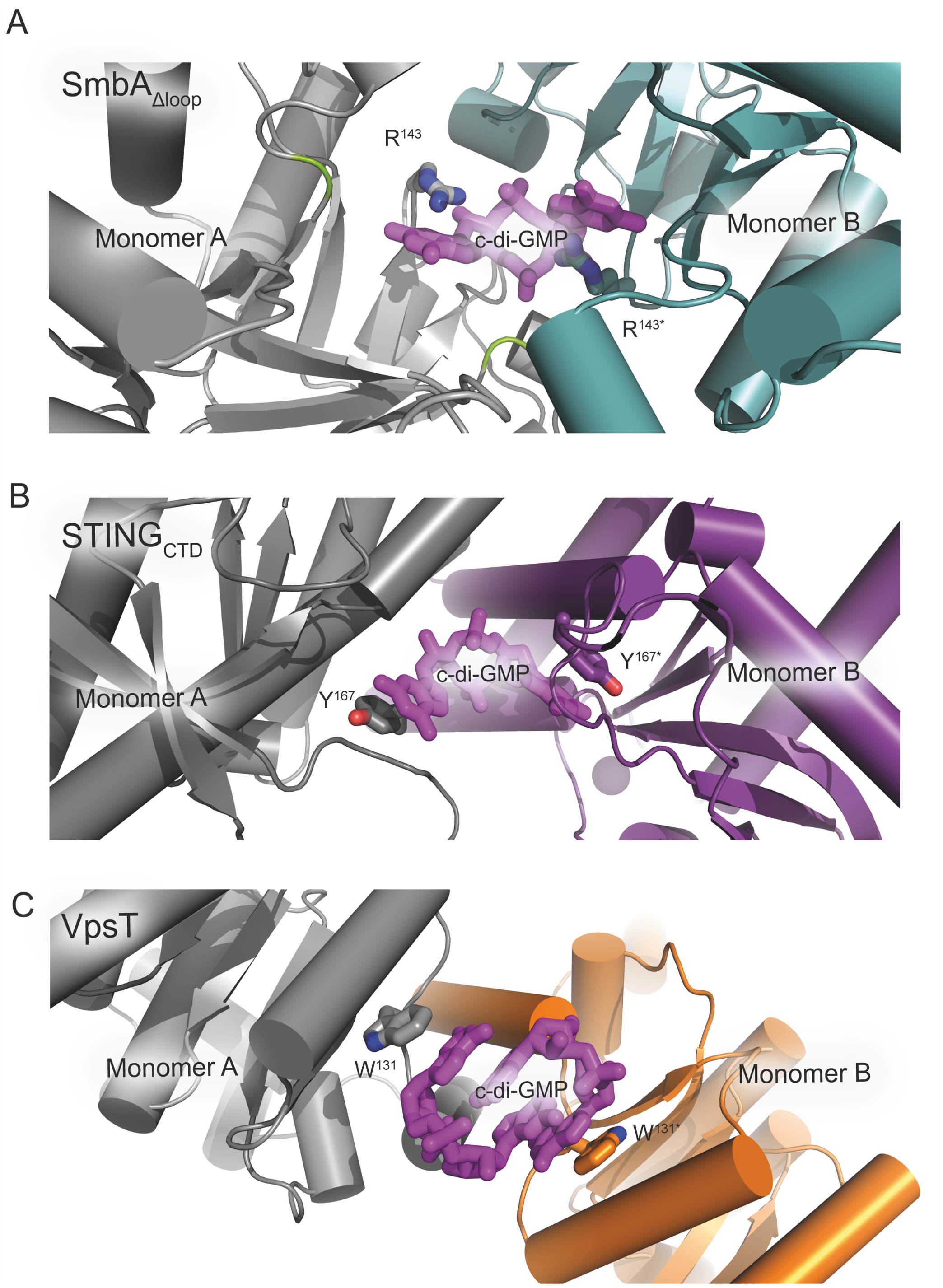


**S2 Fig. c-di-GMP bound at dimer interface found in STING and VpsT.** c-di-GMP bind at the crystallographic dimer interface shown in the stick model (magenta). Residues involved in base stacking are shown in a stick. Monomers are colored differently.

(A) SmbA$_{\Delta loop}$ c-di-GMP contact at the 2-fold crystallographic dimer interface,
(B) STING dimerization interface (PDB code-4F5Y).
(C) VpsT dimerization interface (PDB code-3KLO).

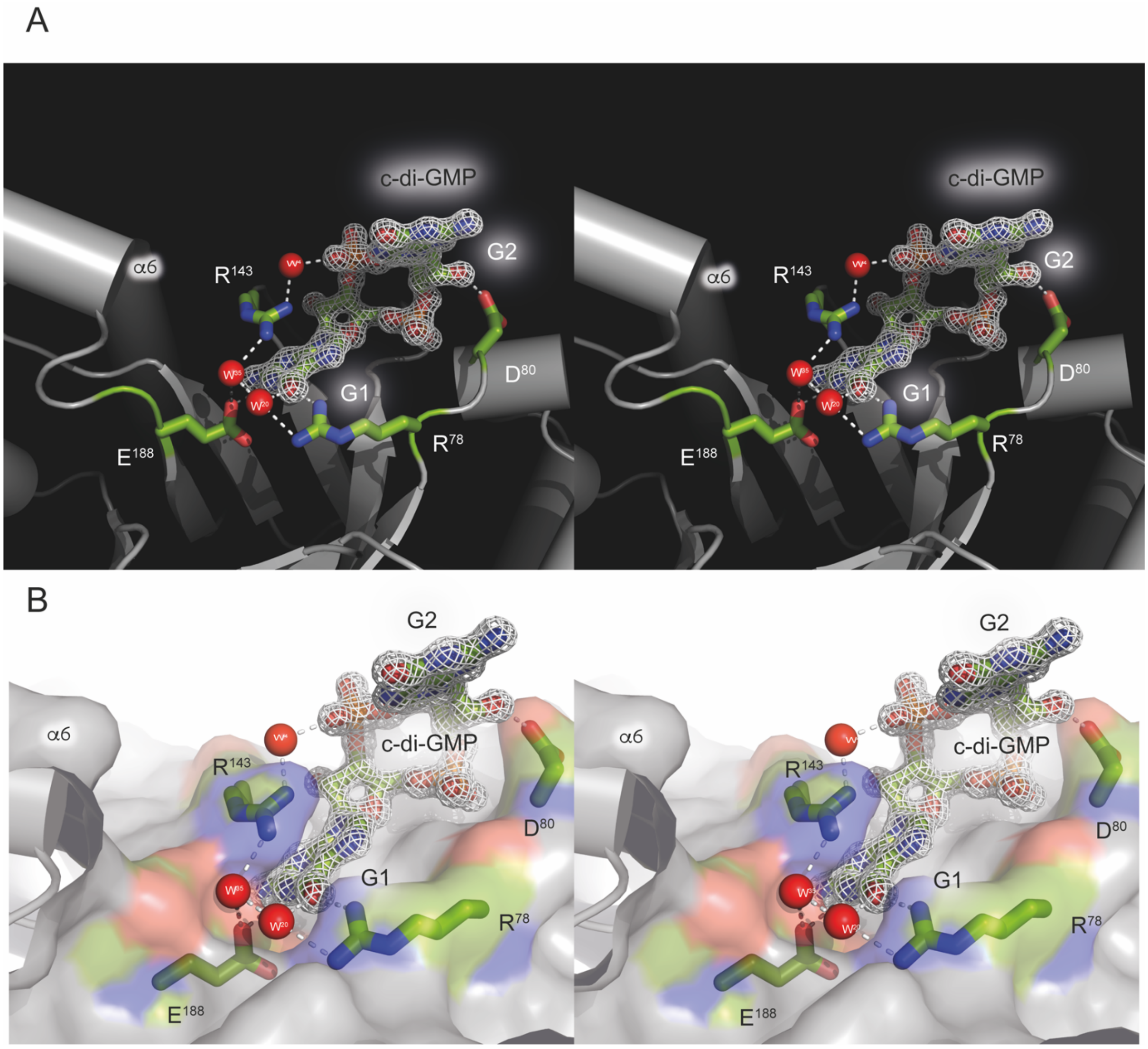


**S3 Fig. An expanded stereo view of the SmbA$_{\Delta loop}$ residues interacting with c-di-GMP.**

**(A)** Stereo view of the 2Fo-Fc omit maps contoured at 1.2 σ. The molecular structure of c-di-GMP is embedded in the map. The colour code is similar to that in Fig. 3A. The R143 guanidinium group stacks very well with the guanyl base of c-di-GMP., while R78 is engaged in lateral H-bonding.

**(B)** Stereo view of the c-di-GMP molecule drawn as surface representation (negatively charged atoms in red, positively charged atoms in blue and carbon atoms in green). Hydrogen bonds are marked by dotted lines in white.
